# Male clients of male sex workers in West Africa: A neglected high risk population

**DOI:** 10.1101/536482

**Authors:** Cheick Haïballa Kounta, Luis Sagaon-Teyssier, Pierre-Julien Coulaud, Marion Mora, Gwenaelle Maradan, Michel Bourrelly, Abdoul Aziz keita, Stéphane-Alain Babo Yoro, Camille Anoma, Christian Coulibaly, Elias Ter Tiero Dah, Selom Agbomadji, Ephrem Mensah, Adeline Bernier, Clotilde Couderc, Bintou Dembélé Keita, Christian Laurent, Bruno Spire, the CohMSM Study Group

**Author notes:** CohMSM study group members are listed in the Appendix. **Corresponding author** (CHK).

## Abstract

Research on male clients of male sex workers (MCMSW) has been neglected for a long time globally. We aimed to characterize MCMSW and to identify factors associated with their sexual practices using data from the prospective cohort study CohMSM conducted in Burkina Faso, Côte d’Ivoire, Mali and Togo. Our study focused on HIV-negative men who have sex with other men (MSM) recruited between 06/2015 and 01/2018 by a team of trained peer educators. Scheduled study visits at 6, 12 and 18 months included medical examinations, HIV screening, risk-reduction counselling and face-to-face interviews to collect information on their sociodemographic characteristics, sexual behaviours, and HIV risk-reduction strategies (HIV-RRS). Three stigmatization sub-scores were constructed (experienced, perceived and internalized). Mixed-effects logistic regression was used for data analysis. Of the 280 participants recruited at baseline, 238, 211 and 118, respectively, had a follow-up visit at 6, 12 and 18 months. Over a total of 847 visits, 47 transactional sex (TS) encounters were reported by 38 MCMSW (13.6%). Of the latter, only one participant reported systematic TS (2.6%), 18 (47.4 %) stopped reporting TS after baseline, and 6 (15.8%) reported TS after baseline. Thirteen participants (34.2 %) reported occasional TS. After adjusting for country of study and age, the following self-reported factors were associated with a greater likelihood of being MCMSW: protected anal sex, exclusively insertive anal sex with male sexual partners, avoidance of sex after consuming psychoactive products and experiencing stigmatization (all during the previous 6 months). The majority of MCMSW in this study employed HIV-RRS with male sexual partners, including engaging in protected anal sex, avoidance of sex when consuming psychoactive products, and practising exclusively insertive anal sex.

## Introduction

Men who have sex with other men (MSM), including male sex workers (MSW), are at a much greater risk of acquiring and transmitting sexually transmitted infections (STI), including human immunodeficiency virus (HIV). This is particularly true in Sub-Saharan Africa, where the prevalence of HIV infection is estimated at 36.3% in MSW compared with 17% in the general MSM population (1–7). Despite the vast literature on sex work, most studies to date have focused exclusively on female sex workers (8–12). Although a few studies have focused on MSW and the risk of HIV/STI transmission (13,14,2,15,16), research on male clients of MSW is very scarce (17–20).

Male clients of MSW (MCMSW) are a key group given their potential role as a bridge for HIV transmission to the general population (17,18,21,22). In Africa, a substantial proportion of MCMSW marry women to avoid experiencing the ongoing hostile anti-MSM social environment. In this way, they can continue to have sexual relations (commercial or not) outside of marriage with male partners. Their non-commercial partners (wife and/or steady male partners) are often unaware of their spouse’s/partner’s commercial sex activities, and the fact that they are being put at a higher risk of exposure to HIV (23,24). A few studies have highlighted that a substantial proportion of MSM who pay for sex and/or have been paid for sex (19,25,26) have risky sexual behaviours, especially condomless insertive or receptive anal sex (27,18,17).

In this study, our main hypothesis was that MCMSW have risk factors for HIV infection which are associated with their sexual practices. Characterizing MCMSW is crucial in order to investigate not only whether a particular profile exists or not for this sub-population, but also to have a better understanding about their psychosocial characteristics and their sexual behaviours. This study had two objectives: first, to compare MCMSW with the general MSM population in terms of their sociodemographic and economic characteristics (e.g., age, educational level, employment status, income) and psychosocial characteristics (e.g. self-defined sexual and gender identities, sudden sexual violence by male partners); second, to identify factors associating MCMSW with TS. The study’s findings will be useful for healthcare providers and researchers because they offer the first comprehensive insight into both MCMSW and the HIV and STI exposure factors associated with this sub-population’s sexual practices.

## Materials and Methods

### CohMSM study procedures

In June 2015, a prospective cohort study of MSM was initiated at the premises of four local community-based organisations providing HIV services to MSM in four West African cities: Abidjan (Côte d’Ivoire), Bamako (Mali), Lomé (Togo) and Ouagadougou (Burkina Faso). Its main objectives were to assess both the feasibility and value of providing novel three-monthly preventive global care for MSM in West Africa, in order to help reduce the incidence of HIV in this key population, in their female partners and in the general population. The study did not compare a control group with an exposed group, nor was it based on a clinical trial. Participants were identified and recruited by a team of trained peer educators from these local organisations who approached individuals through a specific MSM network. Eligibility criteria included being at least 18 years old, and reporting at least one anal sexual intercourse (insertive or receptive) with another man in the 3 months preceding study enrolment. Eligible individuals were offered a quarterly preventive global care package including: i) collection of information on health status, STI symptoms and sexual behaviours of individuals, ii) clinical examination, iii) diagnosis of STI and if necessary their treatment, iv) prevention tips tailored to MSM based on risk-reduction counselling, and v) provision of condoms and lubricants. In addition, vaccination against hepatitis B and annual tests for syphilis were proposed. HIV-negative MSM were also offered an HIV test at each quarterly visit. MSM found to be HIV-positive were offered immediate care for their infection, including ARV treatment. Before starting the interview, participants systematically received detailed information about the survey’s objectives and their right to interrupt the interview without justification. At enrolment and follow-up visits, participants completed face-to-face interviews with a research assistant who collected information on their sociodemographic and economic characteristics, HIV risk-reduction strategies, alcohol consumption, drug use and stigmatization. Participants had to provide written informed consent. The study team was very attentive to ensuring anonymity and confidentiality. Ethics approval was obtained from the National Ethics Committees of Mali (N°2015/32/CE/FMPOS), Burkina Faso (N°2015-3-037), Côte d’Ivoire (N°021/MSLS/CNER-dkn) and Togo (N°008/2016/MSPSCAB/SG/DPML/CBRS). The study protocol was designed in accordance with the ethical charter for research in developing countries of French National Agency for Research on AIDS and Viral Hepatitis (ANRS) in France. The ClinicalTrials.gov Identifier is NCT02626286.

### Current study population

Between 06/2015 and 01/2018, 778 participants were enrolled in CohMSM. All HIV-positive participants (n=154), participants receiving benefits for sex (i.e. money, accommodation or gifts) at least once during the follow-up (n=294) and those who did not complete a sociodemographic questionnaire at baseline (n=50) were excluded from the present analysis (Appendix 1). The present analysis therefore focused on the remaining 280 HIV-negative MSM of whom 238, 211 and 118, respectively, had a follow-up visit at 6-and 12-and 18-months.

### Variables

#### Outcome

The outcome of this study was constructed on the basis of the following question: “During the last 6 months, have you been in a situation where you gave money, accommodation or any other benefit in exchange for sex with a man?”. Participants who responded “always” or “sometimes”, in contrast to those who responded “never”, were categorized as MCMSW. This question was asked at all follow-up visits.

#### Explanatory variables

*a) Socio-demographic and economic characteristics:* age was specified as a continuous variable. Dichotomous variables were constructed to indicate whether participants had at least a high-school level of education (=1 vs. < high-school=0), were married or cohabitating (=1 vs. single, divorced or widowed=0), and whether they had a stable housing status (=1 vs. unstable housing status=0). Socio-economic characteristics included employment status (employed=1 vs. unemployed=0), monthly income dichotomised at the median (50 000 Francs de la Communauté Financière en Afrique, approximately US$89.28 in 2017), source of income (work=1 or aid=0) and self-perceived financial situation (comfortable=1 vs. difficult=0).

b) *Sexual characteristics:* a dichotomous variable indicated self-defined gender identity (both a man and a woman=0 vs. man exclusively=1), and a self-defined sexual orientation identity variable indicated whether participants perceived themselves to be bisexual (=0 vs. not bisexual including homosexual, heterosexual=1). A dichotomous avoidance variable was constructed (no=0 vs. yes=1) to indicate whether participants practiced HIV risk-reduction strategies (e.g., avoiding sexual relations when drunk or when consuming other psychoactive products; using antiretroviral drugs to reduce the risk of HIV infection; avoiding anal penetration by seropositive partners or partners of unknown serostatus, etc.) (Appendix 2). Sexual behaviour was recorded using various variables: i) sexual position taken with male partners (exclusively insertive=0 vs. receptive or versatile=1 and not documented=2); ii) condom use with male partners during anal sex (no=0 vs. yes=1), iii) condom use with male partners during oral sex (no=0 vs. yes=1), iv) number of male sexual partners (more than one=1 vs. one=0), v) disagreement about condom use with male partners (no=0 vs. yes=1), vi) group sex with men (no=0 vs. yes=1). The information provided by all these variables concerned the 6 months before the survey. Another variable, entitled “searching for male sexual partners on the internet” (no=0 vs. yes=1), concerned the previous 4 weeks.

c) *Stigmatization during the previous 6 months:* three sub-scores were constructed ranging from 0 to 10. They were based on items taken from previous study (Appendix 3) (28): 1) “experienced stigmatization” (based on 5 items, Cronbach’s alpha=0.58); 2) “perceived stigmatization” (based on 11 items, Cronbach’s alpha=0.70); and 3) “internalized stigmatization” (based on 8 items, Cronbach’s alpha=0.80).

Socio-demographic and economic characteristics were measured at baseline only and were specified in the model as time-fixed variables. In contrast, sexual characteristics and stigmatization variables were measured at each time-point in the follow-up and consequently were specified as time-varying.

### Statistical analysis

Descriptive analysis was conducted to compare baseline socio-demographic and economic characteristics, and sexual behaviours between MSM practising TS (i.e. MCMSW) and those not practising it. Categorical variables were compared between these two groups using Pearson’s chi-squared test (χ2). Continuous variables were compared using Student’s t-test.

Univariate and multivariate analyses were then performed using mixed-effects logistic regression to account for the correlation of repeated data over time. All explanatory variables were first tested in univariate mixed-effects logistic regression. Potential variables for the multivariate model were then selected with a p-value<0.2. The final multivariate model was estimated using a forward procedure, which consisted in placing all candidate variables into the multivariate model for testing one by one, and then retaining those with a p-value < 0.05. Fixed effects for each study country were specified in order to avoid bias arising from differences in sample sizes. All statistical analyses were performed using Stata version 13.0 (Stata Corp, College Station, Texas, USA).

## Results

### Overall sample description

Of the 280 HIV-negative participants in this analysis, 238, 211 and 118 had a follow-up visit at 6-and 12-and 18-months, respectively. Over a total of 847 visits, 47 TS encounters were reported by 38 MCMSW (13.6%). Of the latter, only one participant reported systematic TS (2.6%), 18 (47.4 %) stopped reporting TS after baseline, and 6 (15.8%) reported TS after baseline. Thirteen participants (34.2 %) reported occasional TS.

**Table 1** shows the comparative analysis of baseline individual characteristics between MCMSW and non-MCMSW. The former were significantly older (average age of 28.5 years vs. 25.4 years, p<0.0017). Furthermore, 68.4% (vs. 51.2%) of them were significantly more likely to have an educational level < high-school diploma. A majority of MCMSW (84.2% vs. 76.9 %) were unmarried (single, divorced or widowed) and although 47.4% had an income generating activity, 71.1% reported their financial situation as difficult. Fifty-four percent (52.6%) had a monthly income above the median (50 000 FCFA). Furthermore, MCMSW were significantly more likely (73.7% increased probability) to report work (as opposed to financial aid) as the main source of their income (p=0.015). Despite having work, they were significantly more likely (50% increased probability, p=0.004) to have unstable housing. Almost half of the MCMSW (47.4%) defined themselves as bisexual while 71.1% identified themselves as being men or boys. MCMSW were also significantly more likely to have exclusively insertive anal sex with male partners.

**Table 1:** Comparative analysis of the baseline characteristics of the study sample (n=280)

| Sociodemographic and economic characteristics | Male Clients<br>n=38 (13.6 %)<br>n (%) | No Male Clients<br>n=242 (86.4 %)<br>n (%) | <sup>a</sup>p Value |
| --- | --- | --- | --- |
| Study countries (n=280) |  |  |  |
| Mali | 15 (39.5) | 62 (25.6) | 0.006 |
| Cote d'Ivoire | 4 (10.5) | 65 (26.9) |  |
| Burkina | 5 (13.2) | 66 (27.3) |  |
| Togo | 14 (36.8) | 49 (20.3) |  |
| Age (n=280) |  |  |  |
| Age [mean ± standard deviation] | [28.5±7.7] | [25.4±5.2] | 0.0017 |
| Education level (n=280) |  |  |  |
| ≥ high-school diploma | 12 (31.6) | 118 (48.8) | 0.048 |
| < high-school diploma | 26 (68.4) | 124 (51.2) |  |
| Marital status (n=280) |  |  |  |
| Married or living in a couple | 6 (15.8) | 56 (23.1) | 0.310 |
| Single, Divorced, Widowed | 32 (84.2) | 186 (76.9) |  |
| Had an income generating activity (n=280) |  |  |  |
| No | 20 (52.6) | 153 (63.2) | 0.212 |
| Yes | 18 (47.4) | 89 (36.8) |  |
| Monthly income relative to the median (n=275) |  |  |  |
| ≤median (50 000 Fcfa) | 18 (47.4) | 141 (59.5) | 0.160 |
| >median (50 000 Fcfa) | 20 (52.6) | 96 (40.5) |  |
| Sources of income (n=280) |  |  |  |
| Aid | 10 (26.3) | 115 (47.5) | 0.015 |
| Work | 28 (73.7) | 127 (52.5) |  |
| Financial perception (n=280) |  |  |  |
| Comfortable | 11 (28.9) | 87 (35.9) | 0.400 |
| Difficult | 27 (71.1) | 155 (64.1) |  |
| Stable housing (n=280) |  |  |  |
| Yes | 19 (50.0) | 177 (73.1) | 0.004 |
| No | 19 (50.0) | 65 (26.9) |  |
| Self-defined sexual identity (n=280) |  |  |  |
| Bisexual | 18 (47.4) | 129 (53.3) | 0.496 |
| Not bisexual | 20 (52.6) | 113 (46.7) |  |
| Self-defined gender identity (n=276) |  |  |  |
| Both a man and woman | 11 (28.9) | 89 (37.4) | 0.314 |
| Man or Boy | 27 (71.1) | 149 (62.6) |  |
| Sexual position with male partners in the previous 6 months |  |  |  |
| Receptive or versatile | 15 (39.5) | 140 (57.9) | 0.043 |
| Exclusively insertive | 23 (60.5) | 96 (39.7) |  |
| ND<sup>f</sup> | 0 (0.0) | 6 (2.5) |  |

<sup>a</sup>p Calculated with Pearson's chi-squared test ( $\chi^2$ ) for categorical variables, Student's t-test for continuous variables.

### Factors associating MCMSW with Transactional Sex

Results from the multivariate analysis (**Table 2**) - after adjusting for the four study countries - showed that the probability of being an MCMSW increased by 4.8% per 1-year increase in age [adjusted odds ratio (aOR) and 95% confidence interval (95% CI):1.048 (1.00-1.10)]. Furthermore, the more participants experienced stigmatization, the higher the probability was that they were MCMSW (this increase reaching 92.0%) (aOR [95%CI]:1.920[1.31-2.81]) during the previous 6-months. In terms of HIV-RRS, participants who self-reported that they practised protected anal sex with male partners were twice as likely to be MCMSW (aOR [95%CI]:2.211[1.15-4.24]) than those who did not report HIV-RRS. Participants who self-reported avoiding sex when drunk or when consuming psychoactive products were 8 times more likely to be MCMSW (aOR [95%CI]:8.789[1.15-67.20]). Finally, participants who self-reported that they exclusively practised insertive anal sex with male sexual partners in the previous 6 months were more than twice as likely to be MCMSW (aOR [95%CI]:2.257[1.12-4.53]).

**Table 2:**
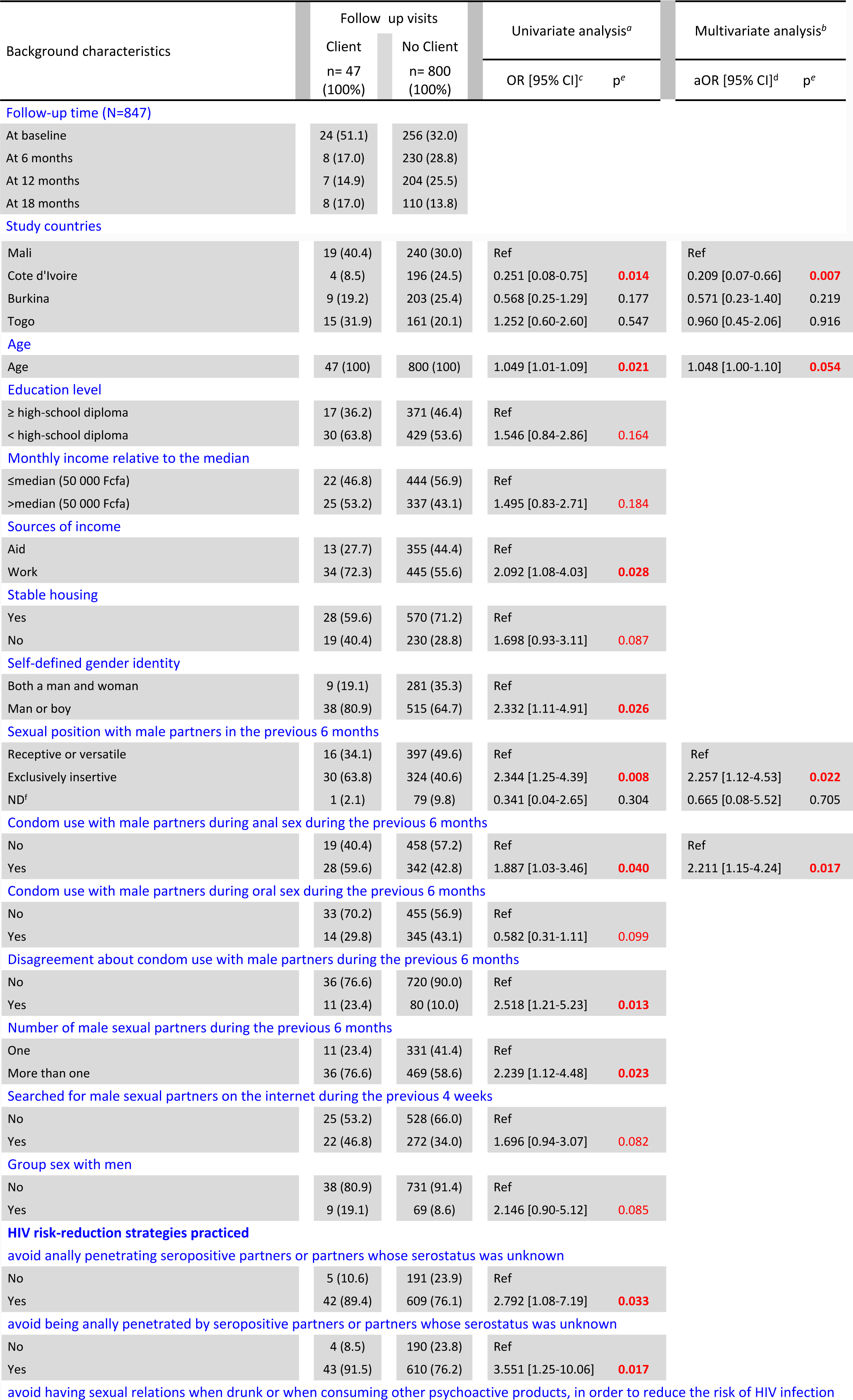
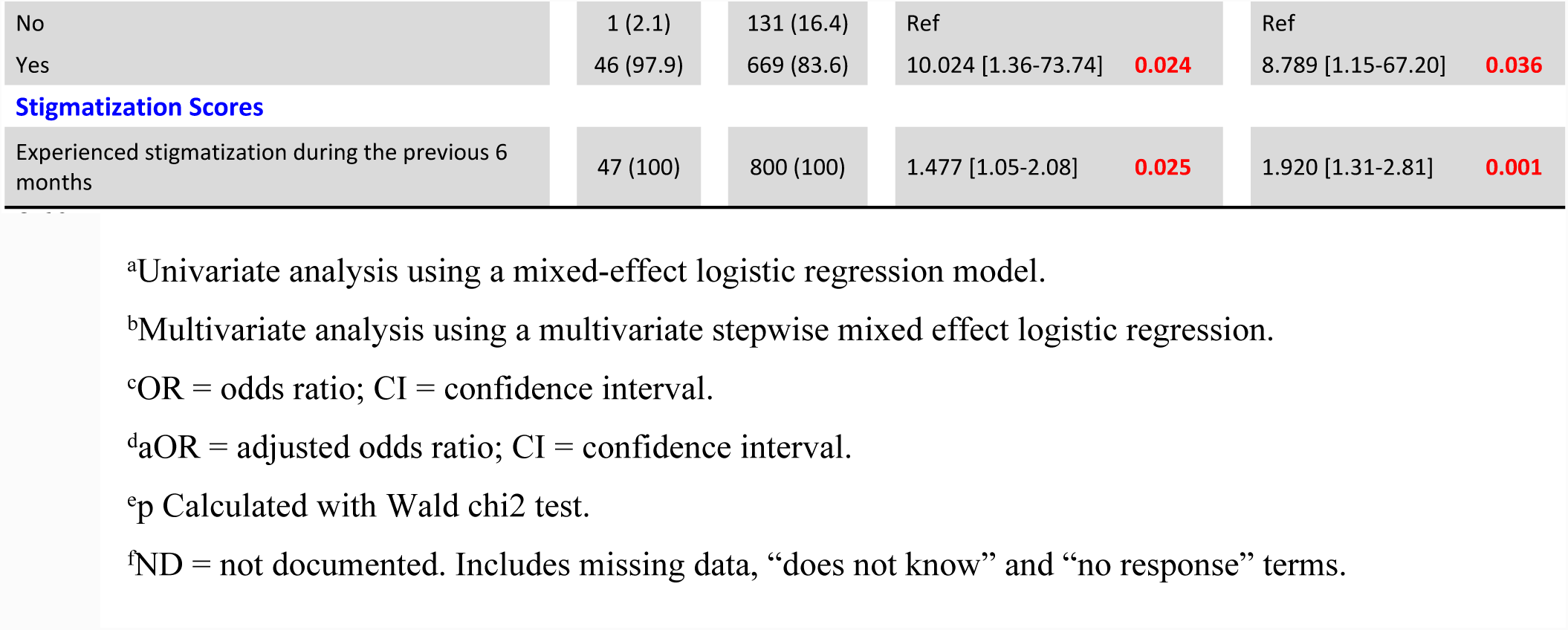
Factors associated with male clients of male sex workers in West Africa: univariate and multivariate analyses with mixed effect logistic regression. (n=280, 847 follow-up visits)

## Discussion

Despite the several studies conducted to date on male clients of Female Sex Workers (29– 32), and the growing amount of literature exploring the experiences and practices of male sex workers (MSW) (14,24,33,34), few studies have focused on male clients of male sex workers (MCMSW). To our knowledge, our study is the first to investigate the issue of transactional sex (TS) in MCMSW in West Africa. Our findings add to the literature by providing information regarding sociodemographic and economic characteristics of this population in four West African countries. About 14 % of our study population were MCMSW (n=38). This percentage is much higher than that found in China (5%) (17). Furthermore, the majority of the MCMSW in our study reported employing risk reduction strategies with male sexual partners, including engaging in protected anal sex, and avoidance of sex when consuming psychoactive products. Similar results for the use of risk reduction strategies among MCMSW were documented in a study in Vietnam in 2013, which first evaluated participants’ perception of risk and then the risk-reduction measures they implemented before sexual intercourse (35). Future intervention efforts should encourage existing HIV risk-reduction practices among MCMSW. Although the subject of a ‘causal relationship’ between homosexuality, unsafe sex and HIV infection has dominated HIV and AIDS discourse in the West (36,37), our findings suggest that pressures, roles and power related to TS may influence the ability of men to behave in ways that reflect their risk perceptions. Accordingly, our results for MCMSW provide some support for health behaviour models which posit that greater perceived risk is associated with fewer risky sexual behaviours. HIV/AIDS education and prevention programs should investigate in greater detail how TS in key populations such as MSM may influence risk perceptions and risk behaviours. Successful HIV prevention strategies for hard-to-reach MCMSW populations require effective integration of evidence-based biomedical, behavioural, and structural interventions, especially in the African context where HIV prevention is centred more on heterosexual contact and vaginal intercourse. As well as social norms, erroneous representations of HIV risks and taboos, HIV prevention programs should also take into account all forms of sexuality and social status (35,38,39). We found that participants who self-reported that they exclusively had insertive anal sex with male sexual partners in the previous 6 months, were more likely to be MCMSW. Previous research showed that insertive anal sex is less risky for HIV contamination than receptive or versatile anal sex (40,41). In our study, the practice of insertive anal sex reported by MCMSW might well contribute to reduce the spread of HIV within and by this population, given that TS is a known factor for increased likelihood of HIV transmission (13). Globally, safer TS plays an important role in risk reduction of HIV and other STI not only for men but also for women (42). This was further indicated by our findings that the probability of being MCMSW tended to increase for each 1-year increase in age. This result contrasts with prior research in the US, China and Australia (19,17,22). The older age of MCMSW might be a barrier to finding younger regular sex partners, which in turn may push them to engage more in transactional sex (43). Finally, our findings also indicated that the more participants experienced stigmatization, the higher the probability was that they were MCMSW. West Africa is hostile in general to homosexuality, but our results do not provide information as to why MCMSW seem to experience more stigmatization than other MSM (24,44–47). The primary strengths of our study come from the fact that the CohMSM study was performed in four West African countries, was longitudinal in nature, and had four scheduled visits. Some study limitations should be taken into account when interpreting our results. First, the CohMSM study was designed to examine the feasibility of introducing a preventive intervention among MSM, and not specifically to explore the issue of MCMSW. Second, given the declarative nature of the data and the fact that respondents participated in face-to-face interviews, social desirability bias is possible. Accordingly, sexual risk behaviours may have been underreported. However, this bias was perhaps minimized by the fact that the research assistants involved all worked close to the ground, came from recognized non-governmental organizations, and were directly involved with the MSM population. It is likely therefore that a trustful relationship emerged over time with the research assistants, and consequently, that they were able to accurately identify MCMSW.

Despite the study’s limitations, our results suggest that MCMSW should be provided with long-term HIV prevention interventions which: (1) focus on individual behaviour change (addressing barriers to condom use with other alternative means of prevention such as pre-exposure prophylaxis (PrEP), enhancing current risk reduction practices); (2) incorporate interpersonal contexts (simultaneously engaging MCMSW and their peers, targeting interpersonal skills, accounting for partner type and intimacy dynamics for regular sexual partners); and (3) take into account their exogenous environments (stigma of being MCMSW in West Africa).

## Conclusions

Little is known about male clients of male sex workers (MCMSW) in West Africa. Our results highlighted a low proportion of MSM who reported gave somethings in exchange for sex (13.6 %) and they were characterized by older age, a lower educational level, an unstable housing and exclusively insertive anal sex. The majority of them employed HIV risk-reduction strategies with male sexual partners, including engaging in protected anal sex, avoidance of sex when consuming psychoactive products, and practising exclusively insertive anal sex.

Despite these positive findings, our study also highlights the need for further research in West Africa targeting MCMSW and their practices, something that has been almost completely neglected to date. A better understanding of this sub-population could help reduce HIV transmission in West Africa.

## Acknowledgements

The study team would like to thank the participants for their valuable contributions to this study. Our thanks also to Jude Sweeney for the English revision and editing of the manuscript.

Contributors
CL and BDK designed and led CohMSM. AAK, SABY, ChC and SA performed data collection under the supervision of CA, ETTD, EM and BDK. GM, MM, MB, and ClC managed the study implementation and field teams. AB trained and supported peer educators.
CHK designed and led the present study under supervision of LST and BS. CHK analysed the data and wrote the first draft of the manuscript. PJC, LST and BS contributed to data analysis, interpretation of the results and correction of the manuscript. All authors critically reviewed and approved the final manuscript.

## References

1. Beyrer C, Baral SD, van Griensven F, Goodreau SM, Chariyalertsak S, Wirtz AL, et al. Global epidemiology of HIV infection in men who have sex with men. The Lancet [Internet]. juill 2012 [cité 20 oct 2016];380(9839):367–77. Disponible sur: http://linkinghub.elsevier.com/retrieve/pii/S0140673612608216

2. Oldenburg CE, Perez-Brumer AG, Reisner SL, Mimiaga MJ. Transactional Sex and the HIV Epidemic Among Men Who have Sex with Men (MSM): Results From a Systematic Review and Meta-analysis. AIDS Behav [Internet]. déc 2015 [cité 15 févr 2017];19(12):2177–83. Disponible sur: http://link.springer.com/10.1007/s10461-015-1010-5

3. Solomon MM, Nureña CR, Tanur JM, Montoya O, Grant RM, McConnell J. Transactional sex and prevalence of STIs: a cross-sectional study of MSM and transwomen screened for an HIV prevention trial. Int J STD AIDS [Internet]. oct 2015 [cité 29 août 2017];26(12):879–86. Disponible sur: http://journals.sagepub.com/doi/10.1177/0956462414562049

4. Baral S, Sifakis F, Cleghorn F, Beyrer C. Elevated risk for HIV infection among men who have sex with men in low-and middle-income countries 2000–2006: a systematic review. PLoS Med [Internet]. 2007 [cité 8 sept 2016];4(12):e339. Disponible sur: http://journals.plos.org/plosmedicine/article?id=10.1371/journal.pmed.0040339

5. Oldenburg CE, Perez-Brumer AG, Reisner SL, Mattie J, Bärnighausen T, Mayer KH, et al. Global Burden of HIV among Men Who Engage in Transactional Sex: A Systematic Review and Meta-Analysis. Prestage G, éditeur. PLoS ONE [Internet]. 28 juill 2014 [cité 13 sept 2016];9(7):e103549. Disponible sur: http://dx.plos.org/10.1371/journal.pone.0103549

6. Smith AD, Tapsoba P, Peshu N, Sanders EJ, Jaffe HW. Men who have sex with men and HIV/AIDS in sub-Saharan Africa. The Lancet [Internet]. 2009 [cité 4 oct 2016];374(9687):416–422. Disponible sur: http://www.sciencedirect.com/science/article/pii/S0140673609611181

7. Stoebenau K, Heise L, Wamoyi J, Bobrova N. Revisiting the understanding of “transactional sex” in sub-Saharan Africa: A review and synthesis of the literature. Soc Sci Med [Internet]. nov 2016 [cité 29 août 2017];168:186–97. Disponible sur: http://linkinghub.elsevier.com/retrieve/pii/S0277953616305305

8. Baral S, Beyrer C, Muessig K, Poteat T, Wirtz AL, Decker MR, et al. Burden of HIV among female sex workers in low-income and middle-income countries: a systematic review and meta-analysis. Lancet Infect Dis [Internet]. 2012 [cité 27 janv 2017];12(7):538–549. Disponible sur: http://www.sciencedirect.com/science/article/pii/S147330991270066X

9. Mountain E, Mishra S, Vickerman P, Pickles M, Gilks C, Boily M-C. Antiretroviral Therapy Uptake, Attrition, Adherence and Outcomes among HIV-Infected Female Sex Workers: A Systematic Review and Meta-Analysis. Sluis-Cremer N, éditeur. PLoS ONE [Internet]. 29 sept 2014 [cité 27 janv 2017];9(9):e105645. Disponible sur: http://dx.plos.org/10.1371/journal.pone.0105645

10. Ngugi EN, Roth E, Mastin T, Nderitu MG, Yasmin S. Female sex workers in Africa: Epidemiology overview, data gaps, ways forward. SAHARA-J J Soc Asp HIVAIDS [Internet]. sept 2012 [cité 27 janv 2017];9(3):148–53. Disponible sur: http://www.tandfonline.com/doi/abs/10.1080/17290376.2012.743825

11. Scheibe A, Drame FM, Shannon K. HIV prevention among female sex workers in Africa. SAHARA-J J Soc Asp HIVAIDS [Internet]. sept 2012 [cité 27 janv 2017];9(3):167–72. Disponible sur: http://www.tandfonline.com/doi/abs/10.1080/17290376.2012.743809

12. Shannon K, Strathdee SA, Goldenberg SM, Duff P, Mwangi P, Rusakova M, et al. Global epidemiology of HIV among female sex workers: influence of structural determinants. The Lancet [Internet]. janv 2015 [cité 27 janv 2017];385(9962):55–71. Disponible sur: http://linkinghub.elsevier.com/retrieve/pii/S0140673614609314

13. Baral SD, Friedman MR, Geibel S, Rebe K, Bozhinov B, Diouf D, et al. Male sex workers: practices, contexts, and vulnerabilities for HIV acquisition and transmission. The Lancet [Internet]. janv 2015 [cité 15 sept 2016];385(9964):260–73. Disponible sur: http://linkinghub.elsevier.com/retrieve/pii/S0140673614608011

14. Mannava P, Geibel S, King’ola N, Temmerman M, Luchters S. Male Sex Workers Who Sell Sex to Men Also Engage in Anal Intercourse with Women: Evidence from Mombasa, Kenya. Cameron DW, éditeur. PLoS ONE [Internet]. 2 janv 2013 [cité 15 sept 2016];8(1):e52547. Disponible sur: http://dx.plos.org/10.1371/journal.pone.0052547

15. Minichiello V, Scott J, Callander D. New Pleasures and Old Dangers: Reinventing Male Sex Work. J Sex Res [Internet]. avr 2013 [cité 23 mars 2018];50(3-4):263–75. Disponible sur: http://www.tandfonline.com/doi/abs/10.1080/00224499.2012.760189

16. Scott J, Minichiello V, Mariño R, Harvey GP, Jamieson M, Browne J. Understanding the New Context of the Male Sex Work Industry. J Interpers Violence [Internet]. mars 2005 [cité 23 mars 2018];20(3):320–42. Disponible sur: http://journals.sagepub.com/doi/10.1177/0886260504270334

17. Chen L, Mahapatra T, Fu G, Huang S, Zheng H, Tucker JD, et al. Male Clients of Male Sex Workers in China: An Ignored High-Risk Population. 1 mars 2016; Volume 71(Number 3). Disponible sur: www.jaids.com

18. Grov C, Wolff M, Smith MD, Koken J, Parsons JT. Male Clients of Male Escorts: Satisfaction, Sexual Behavior, and Demographic Characteristics. J Sex Res [Internet]. oct 2014 [cité 23 mars 2018];51(7):827–37. Disponible sur: http://www.tandfonline.com/doi/abs/10.1080/00224499.2013.789821

19. Grov C, Starks TJ, Wolff M, Smith MD, Koken JA, Parsons JT. Patterns of Client Behavior with Their Most Recent Male Escort: An Application of Latent Class Analysis. Arch Sex Behav [Internet]. mai 2015 [cité 27 mars 2018];44(4):1035–45. Disponible sur: http://link.springer.com/10.1007/s10508-014-0297-z

20. Dizechi S, Brody C, Tuot S, Chhea C, Saphonn V, Yung K, et al. Youth paying for sex: what are the associated factors? Findings from a cross-sectional study in Cambodia. BMC Public Health [Internet]. déc 2018 [cité 24 mai 2018];18(1). Disponible sur: https://bmcpublichealth.biomedcentral.com/articles/10.1186/s12889-017-4999-8

21. Minichiello V, Mariño R, Browne J, Jamieson M, Peterson K, Reuter B, et al. A profile of the clients of male sex workers in three Australian cities. Aust N Z J Public Health [Internet]. oct 1999 [cité 23 mars 2018];23(5):511–8. Disponible sur: http://doi.wiley.com/10.1111/j.1467-842X.1999.tb01308.x

22. Pitts MK, Smith AMA, Grierson J, O’Brien M, Misson S. Who Pays for Sex and Why? An Analysis of Social and Motivational Factors Associated with Male Clients of Sex Workers. Arch Sex Behav [Internet]. août 2004 [cité 23 mars 2018];33(4):353–8. Disponible sur: http://link.springer.com/10.1023/B:ASEB.0000028888.48796.4f

23. Closson EF, Colby DJ, Nguyen T, Cohen SS, Biello K, Mimiaga MJ. The balancing act: Exploring stigma, economic need and disclosure among male sex workers in Ho Chi Minh City, Vietnam. Glob Public Health [Internet]. 21 avr 2015 [cité 15 sept 2016];10(4):520–31. Disponible sur: http://www.tandfonline.com/doi/abs/10.1080/17441692.2014.992452

24. Okal J, Luchters S, Geibel S, Chersich MF, Lango D, Temmerman M. Social context, sexual risk perceptions and stigma: HIV vulnerability among male sex workers in Mombasa, Kenya. Cult Health Sex [Internet]. nov 2009 [cité 14 sept 2016];11(8):811–26. Disponible sur: http://www.tandfonline.com/doi/abs/10.1080/13691050902906488

25. Koken JA, Parsons JT, Severino J, Bimbi DS. Exploring Commercial Sex Encounters in an Urban Community Sample of Gay and Bisexual Men: A Preliminary Report. J Psychol Hum Sex [Internet]. 21 juill 2005 [cité 28 mars 2018];17(1-2):197–213. Disponible sur: http://www.tandfonline.com/doi/abs/10.1300/J056v17n01_12

26. Kumar N, Grov C. Exploring the Occupational Context of Independent Male Escorts Who Seek Male Clients: The Case of Job Success. Am J Mens Health [Internet]. 18 déc 2017 [cité 23 mars 2018];155798831774683. Disponible sur: http://journals.sagepub.com/doi/10.1177/1557988317746836

27. Grov C, Rodríguez-Díaz CE, Jovet-Toledo GG, Parsons JT. Comparing male escorts’ sexual behaviour with their last male client versus non-commercial male partner. Cult Health Sex [Internet]. 7 févr 2015 [cité 23 mars 2018];17(2):194–207. Disponible sur: http://www.tandfonline.com/doi/abs/10.1080/13691058.2014.961035

28. Ha H, Ross MW, Risser JMH, Nguyen HTM. Measurement of Stigma in Men Who Have Sex with Men in Hanoi, Vietnam: Assessment of a Homosexuality-Related Stigma Scale. J Sex Transm Dis [Internet]. 2013 [cité 3 mars 2017];2013:1–9. Disponible sur: http://www.hindawi.com/journals/jstd/2013/174506/

29. Alary M, Lowndes CM. The central role of clients of female sex workers in the dynamics of heterosexual HIV transmission in sub-Saharan Africa. Aids. 2004;18(6):945–947.

30. Jin X, Smith K, Chen RY, Ding G, Yao Y, Wang H, et al. HIV prevalence and risk behaviors among male clients of female sex workers in Yunnan, China. J Acquir Immune Defic Syndr 1999. 2010;53(1):131.

31. Patterson TL, Volkmann T, Gallardo M, Goldenberg S, Lozada R, Semple SJ, et al. Identifying the HIV transmission bridge: which men are having unsafe sex with female sex workers and with their own wives or steady partners? J Acquir Immune Defic Syndr 1999. 2012;60(4):414.

32. Vuylsteke BL, Ghys PD, Traoré M, Konan Y, Mah-Bi G, Maurice C, et al. HIV prevalence and risk behavior among clients of female sex workers in Abidjan, Cote d’Ivoire. Aids. 2003;17(11):1691–1694.

33. Vuylsteke B, Semde G, Sika L, Crucitti T, Ettiegne Traore V, Buve A, et al. High prevalence of HIV and sexually transmitted infections among male sex workers in Abidjan, Côte d’Ivoire: need for services tailored to their needs. Sex Transm Infect [Internet]. juin 2012 [cité 10 sept 2016];88(4):288–93. Disponible sur: http://sti.bmj.com/lookup/doi/10.1136/sextrans-2011-050276

34. Yu G, Clatts MC, Goldsamt LA, Giang LM. Substance use among male sex workers in Vietnam: Prevalence, onset, and interactions with sexual risk. Int J Drug Policy [Internet]. mai 2015 [cité 19 févr 2018];26(5):516–21. Disponible sur: http://linkinghub.elsevier.com/retrieve/pii/S0955395914002928

35. Mimiaga MJ, Reisner SL, Closson EF, Perry N, Perkovich B, Nguyen T, et al. Self-perceived HIV risk and the use of risk reduction strategies among men who engage in transactional sex with other men in Ho Chi Minh City, Vietnam. AIDS Care [Internet]. août 2013 [cité 15 févr 2017];25(8):1039–44. Disponible sur: http://www.tandfonline.com/doi/abs/10.1080/09540121.2012.748873

36. Wolitski, R, R. Valdiserri, P. Denning, W.C Levine. Are we headed for a resurgence of the HIV epidemic among men who have sex with men? Am J Public Health [Internet]. juin 2001 [cité 7 juin 2018];91(6):883–8. Disponible sur: http://ajph.aphapublications.org/doi/10.2105/AJPH.91.6.883

37. CDC, 2005. CDC. 2005. HIV/AIDS Surveillance Report. CDC. http://www.cdc.gov/hiv/topics/surveillance/resources/reports/. :54.

38. Baral S, Logie CH, Grosso A, Wirtz AL, Beyrer C. Modified social ecological model: a tool to guide the assessment of the risks and risk contexts of HIV epidemics. BMC Public Health [Internet]. 2013 [cité 15 févr 2017];13(1):482. Disponible sur: https://bmcpublichealth.biomedcentral.com/articles/10.1186/1471-2458-13-482

39. Möller LM, Stolte IG, Geskus RB, Okuku HS, Wahome E, Price MA, et al. Changes in sexual risk behavior among MSM participating in a research cohort in coastal Kenya: AIDS [Internet]. déc 2015 [cité 27 oct 2016];29:S211–9. Disponible sur: http://content.wkhealth.com/linkback/openurl?sid=WKPTLP:landingpage&an=00002030-201512003-00003

40. Meng X, Zou H, Fan S, Zheng B, Zhang L, Dai X, et al. Relative Risk for HIV Infection Among Men Who Have Sex with Men Engaging in Different Roles in Anal Sex: A Systematic Review and Meta-analysis on Global Data. AIDS Behav [Internet]. mai 2015 [cité 8 juin 2018];19(5):882–9. Disponible sur: http://link.springer.com/10.1007/s10461-014-0921-x

41. Lyons A, Pitts M, Smith G, Grierson J, Smith A, McNally S, et al. Versatility and HIV Vulnerability: Investigating the Proportion of Australian Gay Men Having Both Insertive and Receptive Anal Intercourse. J Sex Med [Internet]. août 2011 [cité 8 juin 2018];8(8):2164–71. Disponible sur: http://linkinghub.elsevier.com/retrieve/pii/S1743609515336183

42. Smith MD, Seal DW. Motivational Influences on the Safer Sex Behavior of Agency-based Male Sex Workers. Arch Sex Behav [Internet]. oct 2008 [cité 23 mars 2018];37(5):845–53. Disponible sur: http://link.springer.com/10.1007/s10508-008-9341-1

43. Bui TC, Nyoni JE, Ross MW, Mbwambo J, Markham CM, McCurdy SA. Sexual Motivation, Sexual Transactions and Sexual Risk Behaviors in Men who have Sex with Men in Dar es Salaam, Tanzania. AIDS Behav [Internet]. déc 2014 [cité 26 sept 2016];18(12):2432–41. Disponible sur: http://link.springer.com/10.1007/s10461-014-0808-x

44. Anderson AM, Ross MW, Nyoni JE, McCurdy SA. High prevalence of stigma-related abuse among a sample of men who have sex with men in Tanzania: implications for HIV prevention. AIDS Care [Internet]. 2 janv 2015 [cité 14 févr 2017];27(1):63–70. Disponible sur: http://www.tandfonline.com/doi/abs/10.1080/09540121.2014.951597

45. Crowell TA, Keshinro B, Baral SD, Schwartz SR, Stahlman S, Nowak RG, et al. Stigma, access to healthcare, and HIV risks among men who sell sex to men in Nigeria. Journal of the International AIDS Society [Internet]. 20:21489. 2017; Disponible sur: http://dx.doi.org/10.7448/IAS.20.1.21489

46. Fay H, Baral SD, Trapence G, Motimedi F, Umar E, Iipinge S, et al. Stigma, Health Care Access, and HIV Knowledge Among Men Who Have Sex With Men in Malawi, Namibia, and Botswana. AIDS Behav [Internet]. août 2011 [cité 15 févr 2017];15(6):1088–97. Disponible sur: http://link.springer.com/10.1007/s10461-010-9861-2

47. Stahlman S, Sanchez TH, Sullivan PS, Ketende S, Lyons C, Charurat ME, et al. The Prevalence of Sexual Behavior Stigma Affecting Gay Men and Other Men Who Have Sex with Men Across Sub-Saharan Africa and in the United States. JMIR Public Health Surveill [Internet]. 26 juill 2016 [cité 29 sept 2016];2(2):e35. Disponible sur: http://publichealth.jmir.org/2016/2/e35/

